# How big is enough? Movement-informed zoning for African swine fever mitigation

**DOI:** 10.64898/2025.12.12.693251

**Authors:** Elodie Wielgus, Tsviatko Alexandrov, Marco Apollonio, Janosch Arnold, Grzegorz Bas, Eric Baubet, Simon Chamaillé-Jammes, Attila Farkas, Claude Fischer, Klemen Jerina, Milos Jezek, Petter Kjellander, Alisa Klamm, Stephanie Kramer-Schadt, Alain Licoppe, Kevin Morelle, Andras Nahlik, Thomas Scheppers, Stefan Suter, Joaquin Vicente, Maik Henrich, Tobias Kürschner, Marco Heurich

## Abstract

1. African swine fever (ASF) poses a serious threat to domestic pigs and wild boar populations. Wild boar can disperse the virus, making effective containment crucial. One of the main control strategies involves establishing restricted zones around detected cases, i.e., areas with temporary restrictions on access and hunting; however, determining the appropriate size of these zones remains a major challenge.
2. To inform the size of restricted zones, we analyzed GPS data from 527 wild boar across 46 European study sites using a two-step approach combining first-passage time analysis and survival modelling to quantify the risk of wild boar leaving areas of different radii (i.e., spatial scales). We investigated how the risk of leaving varied over time and across environmental gradients. To go further, we used our model findings to develop an online application that generates predictive maps of optimal buffer sizes for ASF management at the European scale, based on a given risk threshold (the maximum acceptable probability that a wild boar leaves the area).
3. We found that the relationship between radius and the risk of leaving is negative exponential, and the risk of leaving increased over time, with a more rapid increase for smaller radii. Landscape homogeneity, terrain ruggedness and human impact increased the risk of leaving, with stronger effects at small scales. Contrary to other predictors, agricultural cover exerted a strong effect on risk of leaving over large spatial scales, especially when it was abundant.
4. Across Europe, a buffer radius of ∼8 km is likely sufficient around high-risk infection zones in most areas (considering an infectious period of 14 days and a risk threshold of 5%); however, in certain areas, a radius of up to 20 km may be needed to effectively limit wild boar movement.
5. *Synthesis and applications*: Our results highlight the need for adaptive, context-specific restricted zones. Buffers of 8 km around ASF-affected areas can limit the risk of infected wild boar dispersal, but they may be reduced to 5 km in highly heterogeneous landscapes or high-human impacted areas. Larger buffers may be required in agricultural landscapes. We provide spatially explicit outputs (optimal buffer sizes) that can directly inform policy and wildlife disease response strategies. The approach can be adapted to any other infectious disease.

## Introduction

Wildlife infectious diseases pose significant threats to animal populations and human health. They can result in widespread individual suffering and elevated mortality, population declines, and hence ecosystem disruptions, ultimately leading to biodiversity loss (Smith, Acevedo-Whitehouse & Pedersen 2009). Furthermore, wildlife infectious diseases have the potential to spill over to domestic animal populations and present zoonotic risks to humans (Bengis, Kock & Fischer 2002; Bengis *et al*. 2004), as evident in outbreaks such as Ebola and COVID-19 (Marí Saéz *et al*. 2015; Mazinani & Rude 2021). In recent decades, human population growth, changes in land use inducing habitat fragmentation, and climate change have further increased the risk of disease transmission among and within wild and domestic animal populations as well as humans (Gilchrist *et al*. 2007; Vora 2008). Investing in surveillance, research, and management strategies to prevent and control infectious diseases in wildlife is therefore crucial for safeguarding animal, human, and ecosystem health (Gittleman 2024).

Wild boar *Sus scrofa* are the main wildlife host of African swine fever (ASF) in Europe and are primarily responsible for the transmission of ASF to livestock (Blome, Gabriel & Beer 2013). Since genotype II broke out in 2007 in Georgia, ASF has continued to spread and become more prevalent, raising serious economic, ecological, and animal health concerns across the continent and beyond (Blome *et al*. 2012; Dixon *et al*. 2020). The virus is highly resilient in the environment and in carcasses, and there is no effective vaccine or treatment currently available, making prevention and containment essential to its management. Transmission occurs through direct contact with infected individuals or carcasses, as well as indirectly, as the virus can survive in the environment (e.g., in soil or water) for several weeks (Carlson *et al*. 2020; Mazur-Panasiuk & Woźniakowski 2020; Pepin *et al*. 2020). Anthropogenic transports of fomites or contaminated meat may contribute to long-distance transmission of ASF (Kim *et al*. 2019). Understanding transmission dynamics in the wild remains challenging due to the limited information available on animal contact rates and virus detection in the environment. When considering only non-human-mediated ASF transmission, wild boar movements are presumed to be the main driver of ASF spatial spread, primarily on a local scale due to the severity of the disease (Podgórski & Smietanka 2018). However, there is limited understanding of how environmental and landscape features shape wild boar movement at broad- and disease-relevant scales (Kay *et al*. 2017, K. Morelle, unpublished data, E. Wielgus, unpublished data), such as the critical 14-day window during which most ASF-infected wild boar remain infectious before dying (Gabriel *et al*. 2011; Blome *et al*. 2012; Pietschmann *et al*. 2015), despite the critical importance of this scale for containment efforts.

The current strategy for managing ASF in wild boar populations consists of establishing restricted zones around confirmed cases. These zones are usually designated as follows: an infected zone (all reported cases plus a surrounding buffer where infected animals are likely to move, no disturbances), a buffer zone (minimal disturbance, strict access restrictions, intensive carcass removal), and a control zone (drastic reduction of wild boar densities through intensive culling, Dixon *et al*. 2020; Jori *et al*. 2021; Guberti *et al*. 2022). ASF restricted zones are often defined without clear scientific guidelines, relying on expert opinion or local wild boar monitoring data (if available, Lange 2015; Guberti *et al*. 2019), alongside consideration of available human, logistical, and financial resources. Additionally, decisions are frequently made arbitrarily or uniformly across environmental conditions, overlooking the species’ behavioral plasticity and local context (Podgórski *et al*. 2013; Thurfjell, Spong & Ericsson 2014; D’Amico *et al*. 2015; Stillfried *et al*. 2017; Amendolia *et al*. 2019). Without accounting for wild boar plasticity, restricted zone sizes may be insufficient in some areas or unnecessarily large in others, leading to ineffective and costly disease management. If the restricted zone is too large, the effectiveness of the mitigation measures will be compromised (for example, animal removal may become too labor-intensive to eliminate enough infected animals) and stakeholders, such as farmers, foresters and hunters will be negatively affected. In contrast, if the zone is too small, there is an increased risk of infected animals leaving or have already left the area and spreading the disease elsewhere.

To create a more realistic, species-based zoning system, we used GPS data from healthy wild boar across Europe to quantify the risk that wild boar leave an area of a given size once entered (Freitas *et al*. 2008; Wielgus *et al*. 2023a). We investigated how this risk varies over time, across spatial scales (circle sizes), and under different environmental conditions. This approach is particularly relevant for ASF management as it links movement behaviour to the measurable probability of leaving an area. We demonstrate how this approach can guide the delineation of infected zones by identifying buffer sizes that minimize the likelihood of wild boar leaving an infection focus during the two-week infection window. This provides a science-based tool to support spatial containment strategies.

## Materials and methods

### Movement data and study sites

We used the EUROBOAR database to analyze the risk of wild boar leaving areas of different sizes and investigated the key factors influencing the variation within and between populations. We focused on both male and female yearlings (1-2 years old) and adults (> 2 years old) monitored with GPS collars. We omitted locations recorded during the 10 day post-capture period and collaring to remove potential bias introduced by the initial capture and handling (Stiegler *et al*. 2024).

We used a two-step approach based on first-passage time (FPT) analysis and Cox proportional hazard (CPH) models (Fauchald & Tveraa 2003) to quantify how likely wild boar are to leave areas of different sizes once they have entered them, rather than relying on a simple distance-based approach (e.g., calculating the probability that an individual moves beyond a given distance). This framework treats movement as a dynamic process, estimating when an individual leaves an area after entering it, rather than only whether it ends up beyond a fixed distance. The CPH model uses full-time-to-event information and handles right-censored data. Our approach also allows flexible risk predictions across multiple spatial scales and time windows, which is particularly relevant for defining ASF buffers.

FPT analysis is most effective when data are collected at regular time intervals (Fauchald & Tveraa 2003), and its accuracy depends on the duration of monitoring (Pinaud 2008). We therefore (1) selected individuals for which GPS-loggers were programmed to record location data every 2 h or less and we resampled each individual’s trajectory to have one fix every 2 h (mean = 11.14 locations per day), (2) divided the individual’s trajectory into segments in which successive locations followed this 2-hour interval with a maximum allowable gap of 12 hours, and (3) retained only segments that lasted at least 24 hours. We allowed a maximum gap of 12 hours because wild boar typically move no more than 1.5 km per day (maximum distance between their two furthest locations, E. Wielgus, unpublished data), making it unlikely that they would leave our smallest analysis radius (1 km; see below) within that time.

The final dataset consisted of location data of 527 wild boar monitored from 2005 to 2022 across 46 study sites in 13 European countries. In total, we analysed 1,059,304 locations grouped into 2,732 segments, with segments lasting on average 34.26 (range: 1-549 days). The study sites cover contrasting environmental conditions and are located along a wide latitudinal gradient (Figure 1).

**Figure 1.**
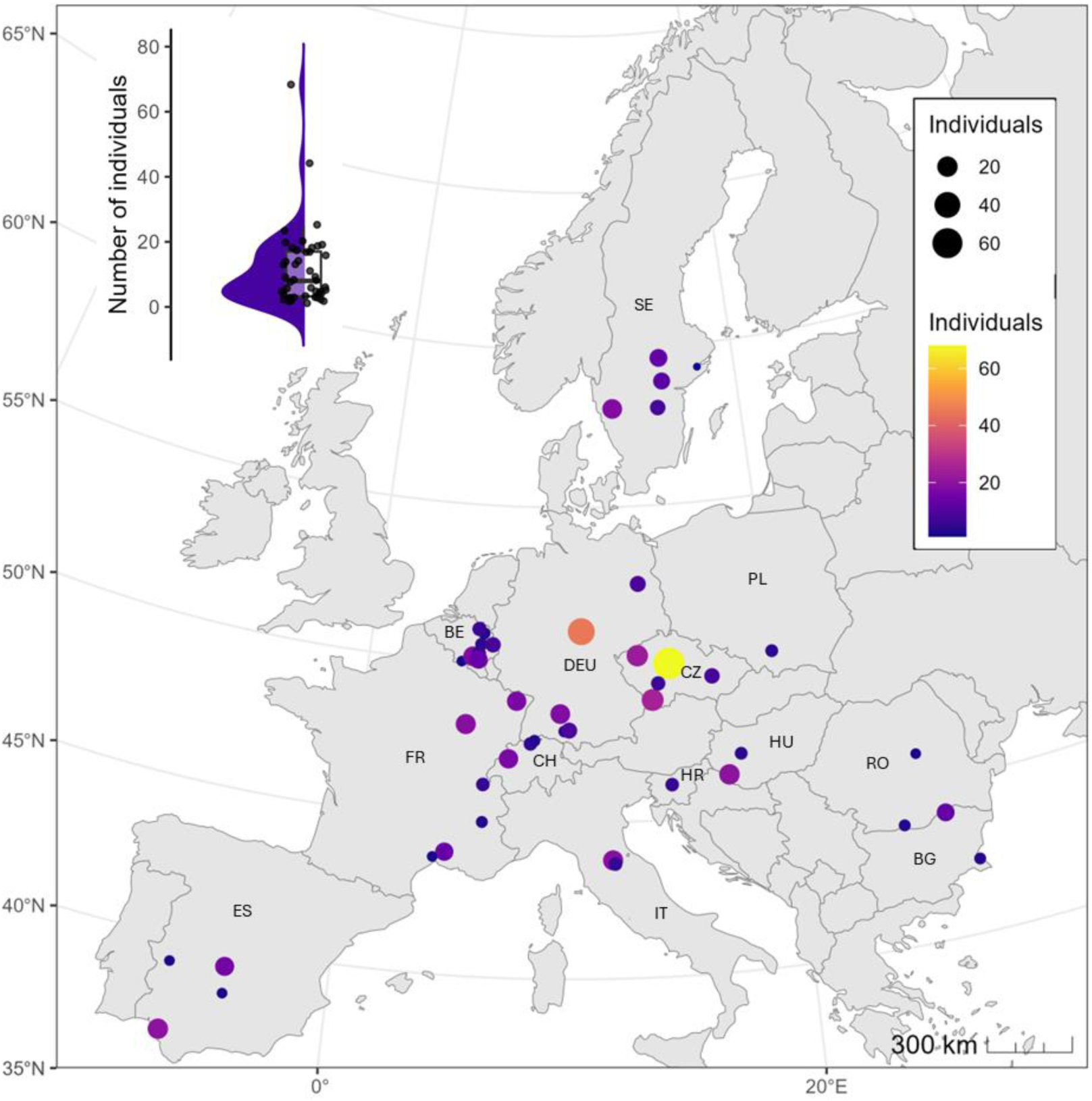
Location of the 46 study sites in Europe. Colour and size of the circle indicate the number of individuals per study site. The small inset at the top left shows the distribution of individuals by study area.

### Residency time assessed using first passage-time

We calculated FPT along each trajectory segment and measured the time an individual spends inside a circle of radius *r* centered on a given GPS location. FPT is a measure of the time spent in the vicinity of a point of interest and can be interpreted more generally as the residency time within a given radius around a focal location (i.e., how long an animal remains within a radius *r* once it has entered it).

To capture how movement behaviour changes with spatial scales, we calculated FPT for different circle sizes, which we refer to as scales. For each trajectory segment, we quantified the residence time within circles of increasing radius (i.e., increasingly large areas) centered on each GPS location. Scales ranged from 1 to 20 km, with 1-km increments between 1 and 10 km and 5-km increments afterwards. When the exact entry and/or exit crossing of the circle was uncertain, we used the first and/or last known location within the circle to estimate the minimum time spent inside. These residency times were then used as input in a CPH model to (1) estimate the probability (risk) that an individual leaves a circle of radius *r* after entering it, and (2) as a function of time and environmental conditions (see *Statistical analyses*). Importantly, this risk corresponds to the likelihood of exiting the area defined by radius *r*, not the probability of moving beyond radius *r* from the focal GPS point. This two-step framework is particularly well-suited for defining restricted zones because it links movement behaviour to a quantified probability of leaving an area, allowing managers to identify the minimum buffer size needed to keep a targeted proportion of individuals contained around an infection point.

### Environmental variables

To investigate the factors that may influence the risk of leaving an area, we extracted a set of explanatory predictor variables describing climate, habitat, topography, and human impact at different spatial scales (Table 1). All the variables were extracted at or resampled to a 100 m resolution.

**Table 1.**
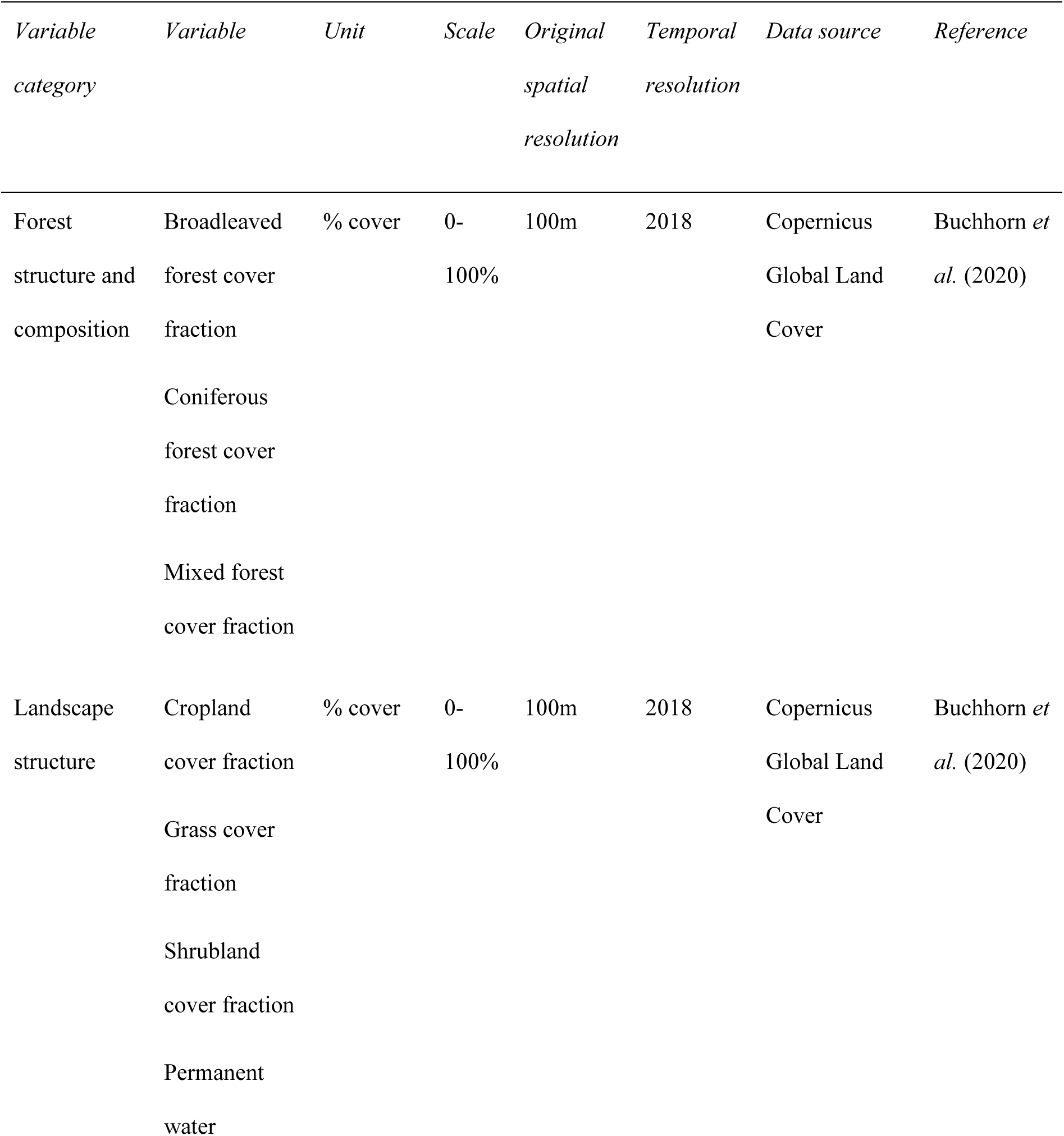
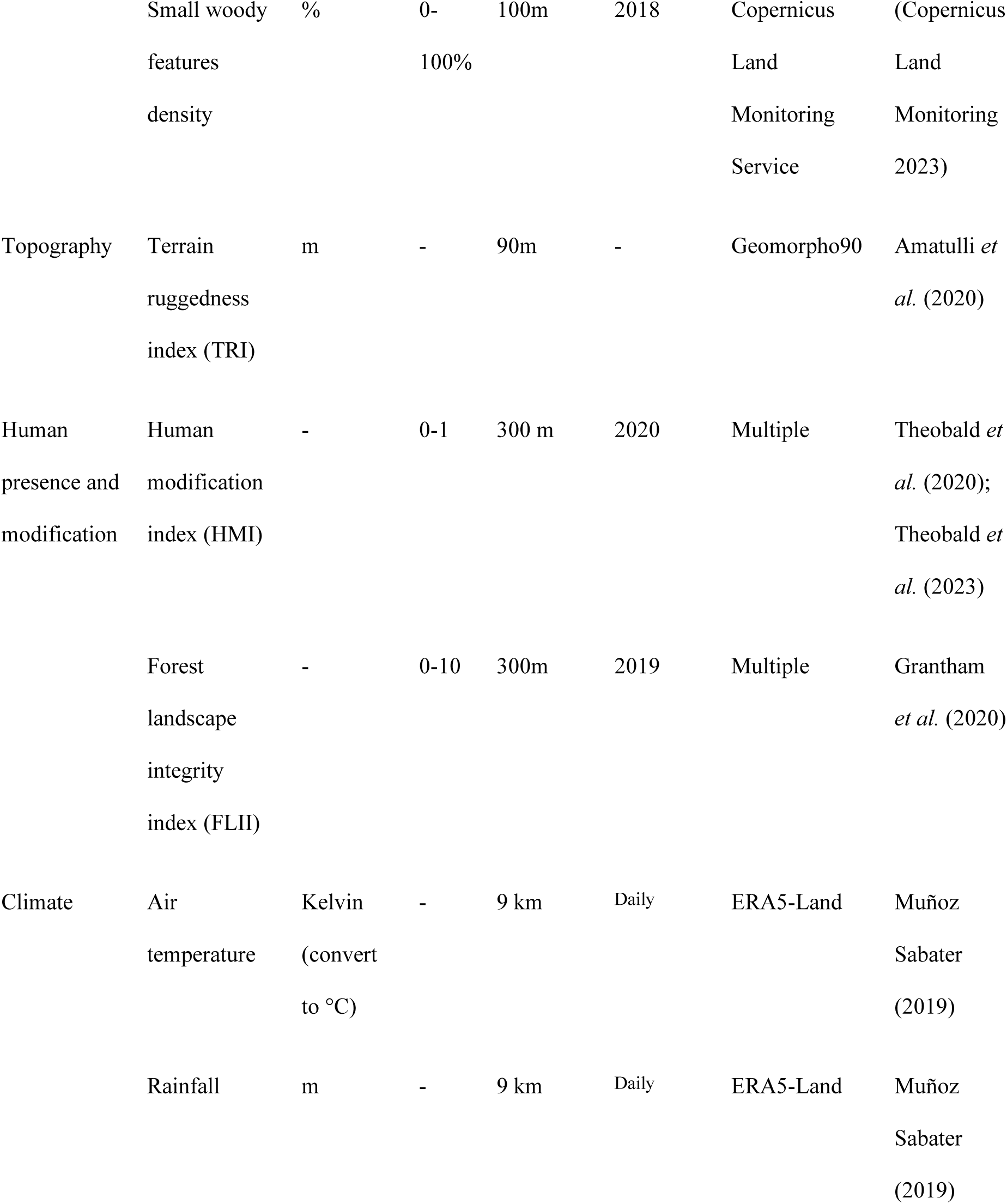
List of environmental predictor variables used to characterize residence times, derived from the first-passage time analysis of wild boar in Europe, including descriptions and sources for each variable.

We described habitat by including shrubland-, cropland-, grass- and permanent water-cover fractions (Buchhorn et al., 2020). To get broadleaved-, coniferous- and mixed-cover-fractions, we used the forest type (Copernicus Land Monitoring 2018) and the forest cover fraction (Buchhorn *et al*. 2020) rasters. Using Quantum GIS, we separated the forest type raster into three distinct layers: Broadleaved forest, Coniferous forest, and Mixed forest, and incorporated the raster cell-level information from the fractional cover raster into each of these new layers. We additionally included small woody features (e.g., hedgerows, tree lines) (Copernicus Land Monitoring 2023) as a separate predictor because they provide cover and foraging opportunities at fine spatial scales and contribute to habitat connectivity (Thurfjell *et al*. 2009; Ferens, Załuski & Borkowski 2025).

To characterize the topographic heterogeneity, we used the terrain ruggedness index available through Geomorpho90, a suite of global high-resolution geomorphometry variables (Amatulli *et al*. 2020).

To characterize the presence of human pressures in the landscape, we included two predictors, the forest landscape integrity index (FLII, Grantham *et al*. 2020), providing a measure of the modification and connectivity of forest areas, and the human modification index (Theobald *et al*. 2020), combining 14 anthropogenic stressors (e.g., human population density, infrastructures, and land use by agriculture) to quantify the degree of land modification.

We described the environmental context of each FPT residency time using a multi-scale moving window. For each spatial scale (circular buffers ranging from 1 to 20 km in radius), we calculated the focal mean values of all predictor variables. These scale-specific averages were then linked to the corresponding FPT residency time. Climatic (temperature and precipitation) values were extracted at the GPS location, assuming negligible variation at a scale of 20 km. We used the values closest in time to each GPS position, at the finest available temporal resolution (daily, see Table 1 for data sources and derivation details.

Extraction of environmental variables was performed using R and Google Earth Engine (https://earthengine.google.com/), a cloud platform for the analysis of geospatial data. All variables were continuous, as land cover was calculated as the proportions of land cover types per grid cell to preserve data quality when rescaling (Seebach *et al*. 2011). Variables were tested for multicollinearity using Spearman’s rank correlation and variables with | *r* | > 0.75 were not included in the same model (Fig S1 Supplementary Information).

### Statistical analyses

We fitted a CPH model to predict the probability (i.e., risk) that a wild boar leaves an area of radius *r* once it has entered it, and to assess how environmental predictors influence this risk. The residency time within each circle, derived from the FPT and representing the time spent inside a radius *r* until the animal exits, was treated as the survival time (the response variable in CPH models). When the exact entry and exit time could not be determined and only a minimum residency time could be estimated, these observations were treated as right censored, reflecting lower confidence in their duration compared to those with known entry and exit times.

As explanatory variables, we included the scale (1-20 km) and all predictors. For predictors extracted at each spatial scale (i.e., all except climate predictors), we included interactions with the scale to capture potential scale-dependent effects. To account for repeated, non-independent observations from the same individuals, we fitted a CPH model with a frailty term for individual identity using the *survival* package (Therneau 2021). The frailty term is a penalized partial likelihood approximation to a random effect.

For each predictor, we calculated hazard ratios (HRs), which represent the relative change in the hazard of leaving associated with a one-unit increase in that predictor, from the exponential of each coefficient (Murray 2006; Freitas *et al*. 2008). In this parameterization, as all predictors are continuous, the reference corresponds to a hypothetical individual with all predictors set to zero. Thus, HRs quantify the multiplicative effect of each predictor relative to this reference. We derived survival functions to estimate how the risk of leaving varies over time and in relation to the predictor variables. We predicted how the risk of leaving changes with the predictor variables for a given time window: a 14-day period, corresponding to the critical timeframe during which most wild boar infected with the highly virulent ASF strain die. All analyses were carried out in the R statistical computing environment (R Core Team 2025).

### Defining buffer for ASF management

To support ASF management, we used the model outputs to predict the optimal buffer size around an infection point (generally, an ASF-positive carcass), accounting for local environmental conditions. Using predictor layers processed through multi-scale moving windows, we generated continent-wide maps predicting the risk that wild boar exits a circular area of radius *r* once it has entered for a given time window (e.g., 14 days). Based on these maps, we then identified, for each cell, the smallest radius for which the predicted risk of leaving remained below a chosen threshold. This threshold represents the maximum acceptable probability that an individual exits the area after entering it. For example, a threshold of 0.05 means that no more than 5% of wild boar will leave the buffer once inside. The smallest radius for which the risk of leaving is less than or equal to this threshold yields the ‘optimal buffer size’ for that location. In other words, this is the distance around a point of infection that 95% of wild boar are expected not to exceed and can thus be used to delineate a restricted zone centered on that point. Under a more permissive threshold (e.g., 0.15), managers would accept that up to 15% of individuals may leave the area. The resulting map, therefore, shows managers the minimum buffer sizes needed to contain wild boar movements within the critical infection window.

To provide greater flexibility for management decision-making, we also developed an interactive application in which users can adjust both the risk threshold (0.001, 0.01, 0.05, 0.1, and 0.15) and the time window of interest (14 or 21 days). The tool then returns the corresponding optimal buffer size needed to contain wild boar movements under conditions at any given location. Users can download the results in GeoTIFF, PNG, or CSV formats, allowing easy integration into mapping workflows and reports. This tool provides a flexible way to adapt management recommendations to local contexts.

We checked whether our model made predictions under environmental conditions that differed from those used for training, as such cases may lack statistical validity and biological relevance. To evaluate this, we determined whether the predictor values of each raster cell fell outside the training range for each spatial scale (radius), using the rasters created by moving windows. We then produced a map showing the average proportion of radii for which this occurred for each raster cell (see Supplementary Information).

To assess the validity of our approach, we compared our predicted optimal buffer sizes with those implemented during past ASF outbreaks in European wild boar populations that were successfully eradicated, specifically in the Czech Republic (2017) and Belgium (2018), in heterogeneous landscapes consisting of forest and rugged terrain. We obtained the locations of 697 wild boar carcasses confirmed to have died from ASF in these two countries from the FAO database. Each carcass was treated as the center of a buffer, and we extracted the predicted optimal buffer size for that location across several risk thresholds (i.e., 0.01 and 0.05) values and time windows (i.e., 14 and 21 days). The 21-day window is also biologically relevant and conservative, as it encompasses approximately one week of incubation followed by a two-week critical period during which most infected individuals die.

## Results

### Survival model outputs

Unsurprisingly, the CPH model revealed that the risk of leaving an area decreased non-linearly with increasing spatial scale at any time, and increased non-linearly with time (Figures 2-3, Table 2). Hazard ratios (HR) are relative to the baseline hazard (i.e., all covariates = 0). At this baseline, the risk of leaving increased with higher proportions of forest (broadleaved, coniferous, and mixed), grassland, shrubland, and water, indicating that more homogeneous landscapes providing cover and shelter are associated with a higher risk. Small woody features and terrain ruggedness further increased the risk. In contrast, human impact and a higher proportion of cropland reduced the risk of leaving. The HR of interaction terms indicates that the effects of the environmental predictors strongly depended on the scale considered (Table 2).

**Figure 2.**
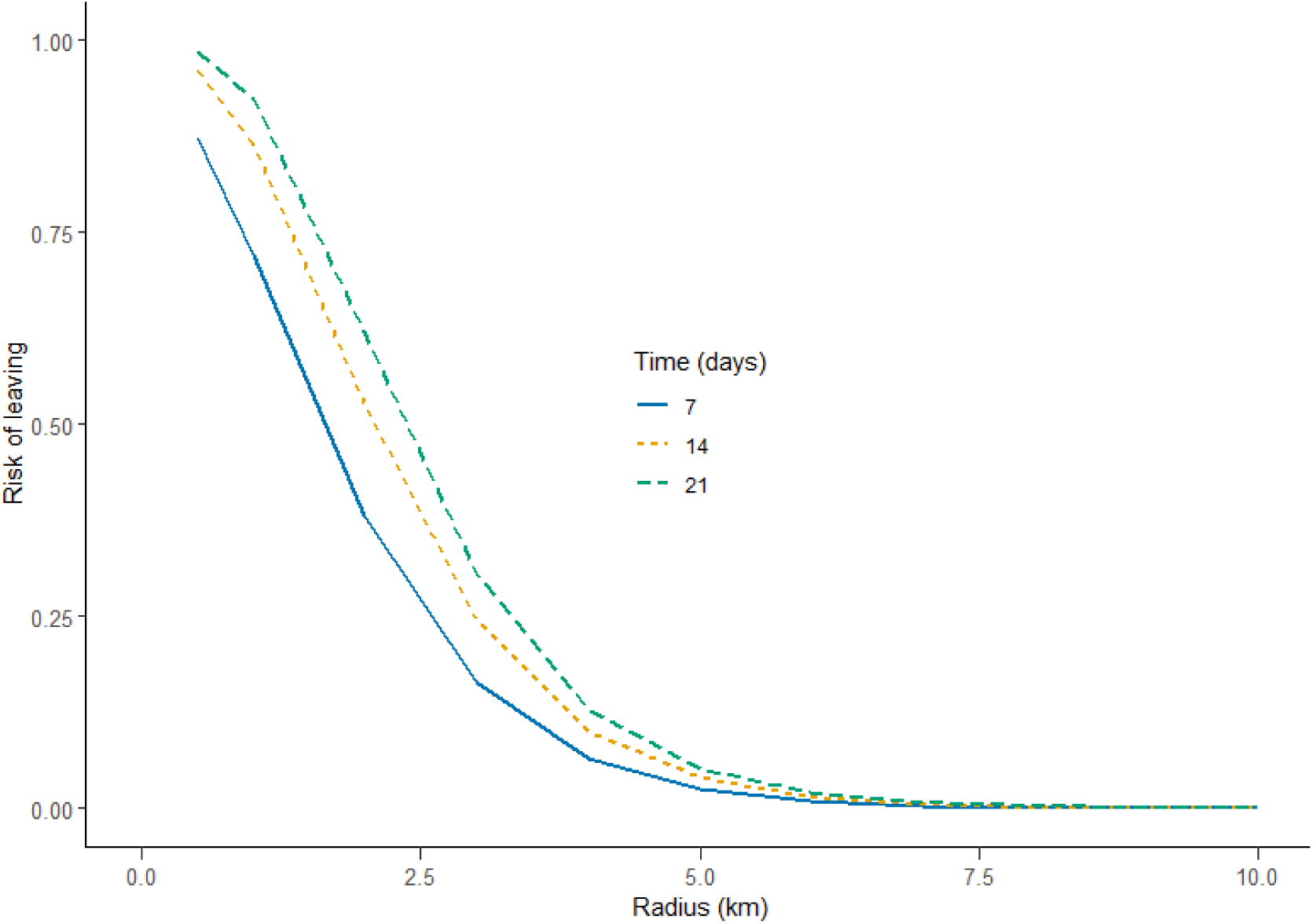
Relationship between the predicted risk of leaving and radius size at 7, 14, and 21 days, based on the Cox proportional hazard model. All other predictor variables were held at their median values.

**Figure 3.**
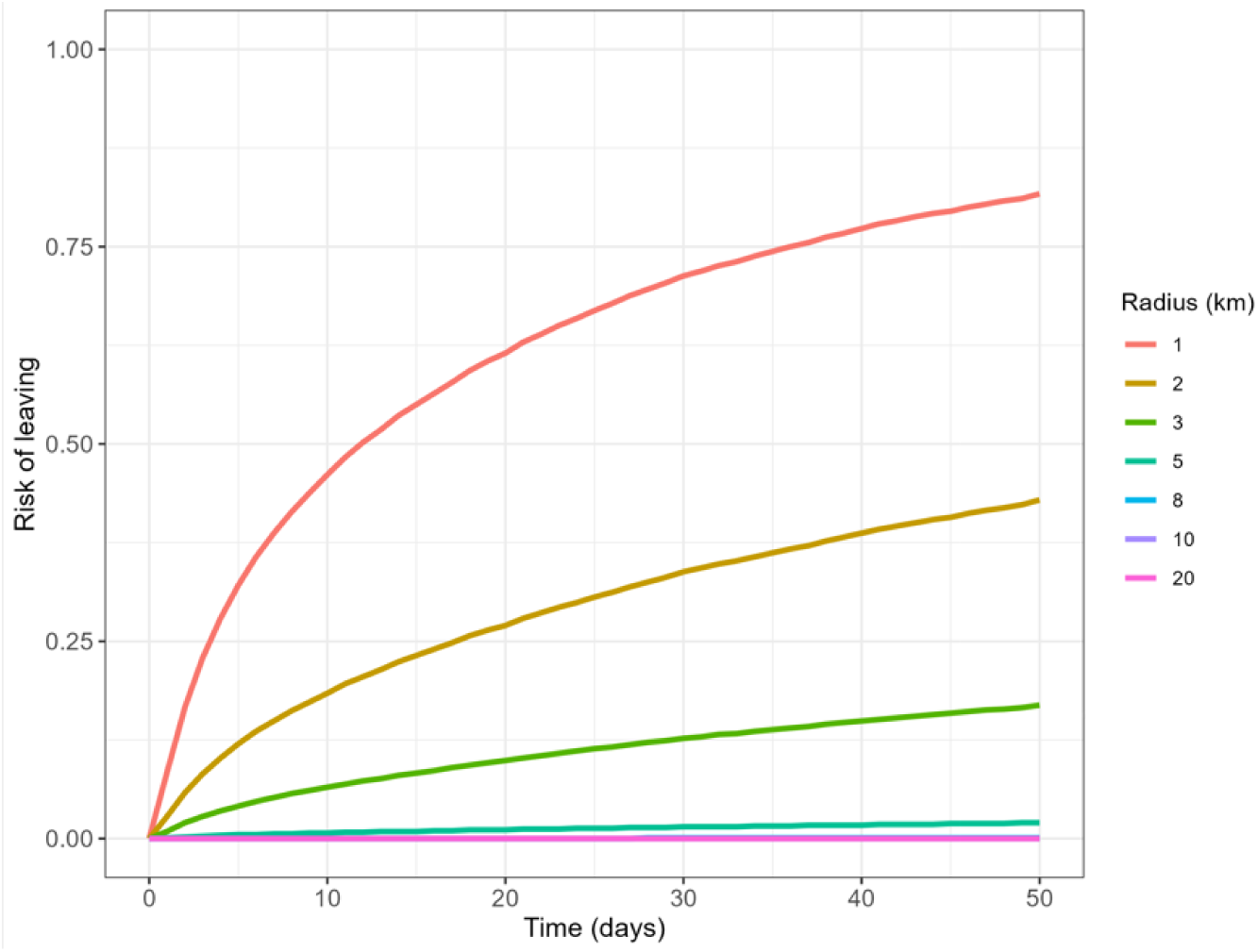
Risk of leaving over time and radius, with all predictors held at their median values per given radius size.

**Table 2.** Estimated coefficients (β) and hazard ratios (e^β^) of the Cox proportional hazard model for the predictor variables, some of which are in interaction with radii reflecting different spatial scales. The variance component attributed to individual variability and the standard deviation of the per-individual random effects (frailty) are also given.

| Variable | HR ($e^{\beta}$) | $\beta$ |
| --- | --- | --- |
| Total precipitation (mm) | 1.001 | 0.001 |
| Temperature | 0.998 | -0.002 |
| radius (km) | 0.104 | -2.268 |
| Broadleaved forest | 1.056 | 0.055 |
| Coniferous forest | 1.025 | 0.025 |
| Cropland | 0.990 | -0.009 |
| Forest integrity index | 1.000 | 0.000 |
| Grass cover | 1.010 | 0.010 |
| Human modification index | 0.764 | -0.269 |
| Mixed forest | 1.055 | 0.054 |
| Permanent water | 1.033 | 0.033 |
| Shrubland | 1.040 | 0.039 |
| Small woody features | 1.052 | 0.050 |
| Terrain ruggedness | 1.010 | 0.010 |
| Radius:Broadleaved forest | 0.988 | -0.012 |
| Radius:Coniferous forest | 1.008 | 0.008 |
| Radius:Cropland | 1.019 | 0.019 |
| Radius:Forest integrity index | 1.000 | 0.000 |
| Radius:Grass cover | 1.020 | 0.019 |
| Radius: Human modification index | 1.569 | 0.450 |
| Radius:Mixed forest | 0.997 | -0.003 |
| Radius:Permanent water | 0.991 | -0.009 |
| Radius:Shrubland | 1.018 | 0.018 |
| Radius:Small woody features | 1.030 | 0.030 |

| Variable | HR ($e^\beta$) | $\beta$ |
| --- | --- | --- |
| Radius:Terrain ruggedness | 1.012 | 0.010 |
| frailty(animals id) | Variance = 1.46 |  |

The variance of individual frailty effects indicates moderate unexplained heterogeneity among individuals (Table 2). The distribution of individual frailty effects (log-hazard) shows that most individuals have risks close to the population average, but a few have substantially lower hazards (Figure S3).

### Temporal and environmental patterns in predicted risk

The predicted risk of leaving at *t* = 14 days increased with all land cover types, including broadleaved, coniferous, and mixed forest, cropland, shrubland, grassland, and water. Small woody features, terrain ruggedness, and human impact further increased the risk of leaving. However, the magnitude of this effect depended on the spatial scale considered. At smaller scales, the effect of these predictors on the risk of leaving was stronger, whereas at larger scales the effect was attenuated, except for cropland. Forest integrity index reduced the predicted risk of leaving when *r* = 1 km, but the direction of the effect slightly changes with increasing *r* to become positive when *r* ≥ 3 km. Most predictors showed only weak or no effect when considered at broader scales (≥ 10 km). Climatic conditions had a negligible effect.

It is important to note that the HRs for cropland and human modification index were both associated with a slightly reduced risk of leaving (HR < 1, Table 2), appearing inconsistent with the predicted curves (Figure 4). This is because HR is for the baseline, i.e., when radius size is equal to 0. However, the interactions between cropland and radius, and human modification index and radius were positive. Once the interaction is accounted for, the overall effects become positive at realistic radius sizes: At scale ≥ 1 km, the interaction term already outweighs the main effects, leading to an increasing risk of leaving with a higher cropland proportion and human impact, as seen on the predicted curves.

**Figure 4.**
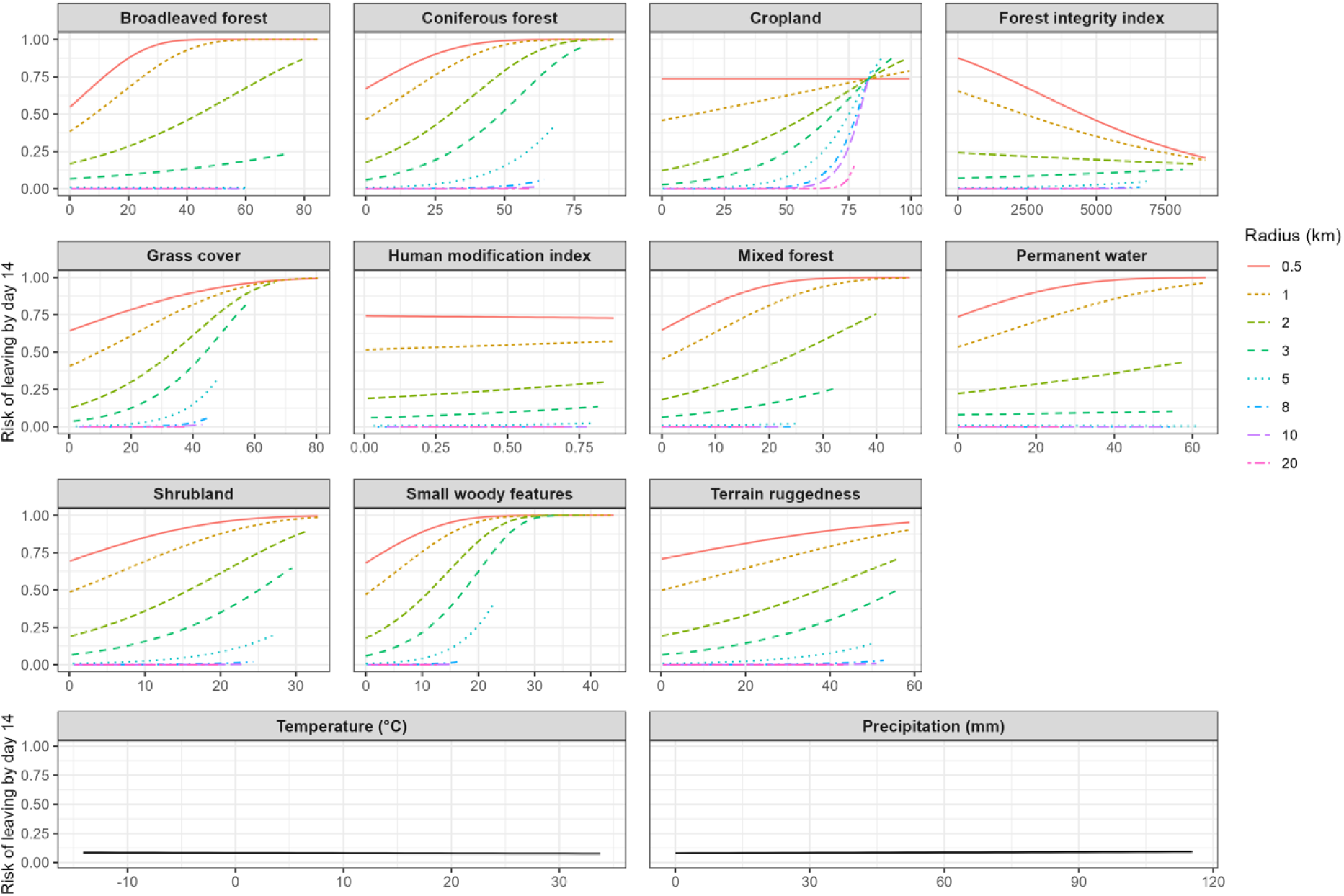
Risk of leaving after 14 days as a function of radius and one varying predictor. Predictions are based on the Cox proportional hazards model, with all other variables held at their median values.

**Figure 5.**
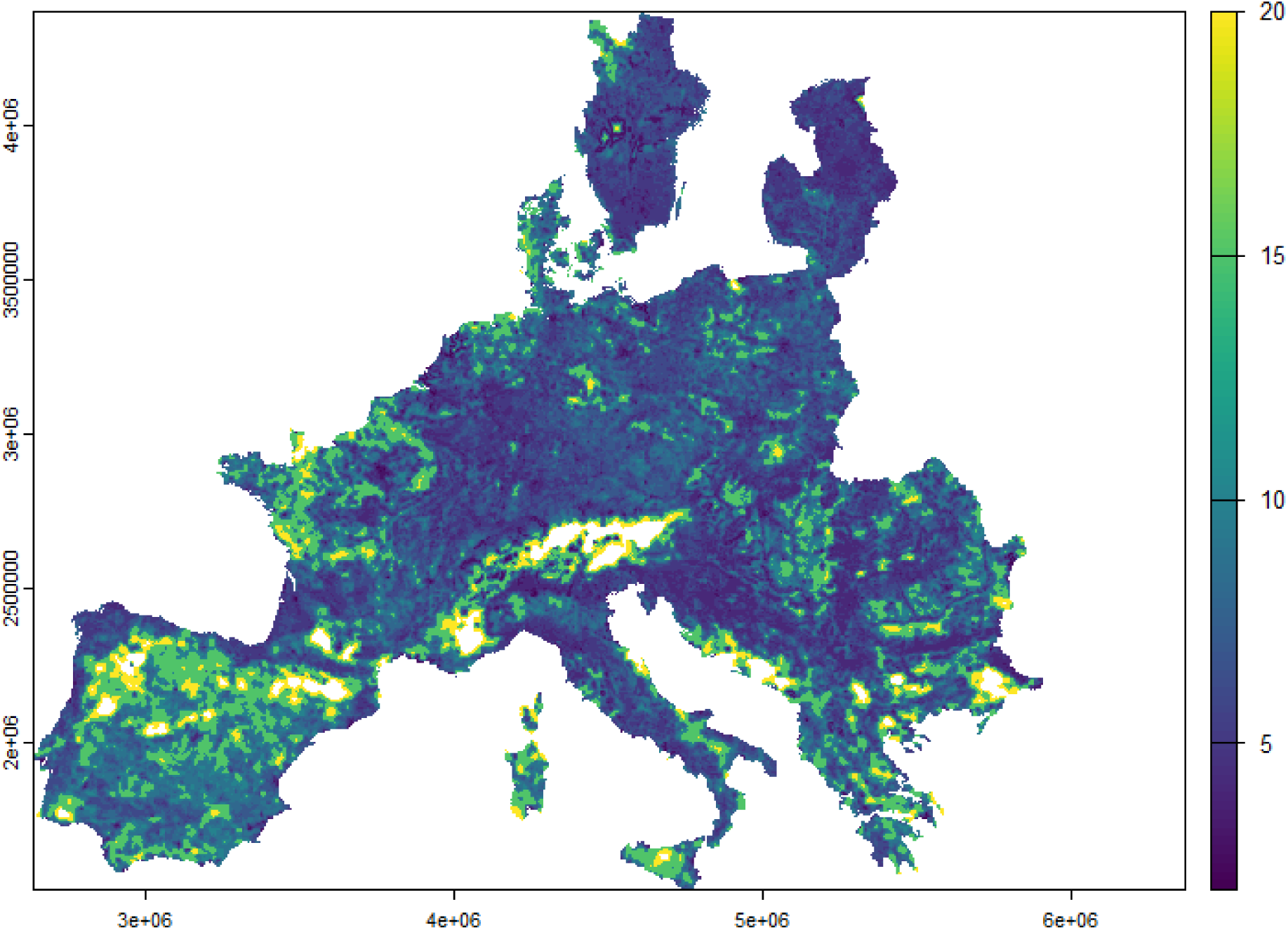
Predicted minimum buffer size required to keep the 14-day risk of wild boar leaving ≤ 0.05 across Europe. Predictions are shown for a temperature of 11°C and precipitation of 0.43 mm, which correspond to the median conditions across our dataset. Map resolution is 100 m. Darker blue areas indicate small buffer radii, yellow areas correspond to larger radii (in km), and white areas indicate radii exceeding 20 km. The map illustrates locally adapted buffer sizes that can guide ASF management.

### Optimal radius size determination

Across Europe, the mean optimal radius size ± standard deviation over a 14-day risk period is 11.01 ± 4.51 km at a 0.01 risk-threshold, 8.23 ± 3.89 km at a 0.05, and 6.89 ± 3.39 km at a 0.10, reflecting the expected decrease in radius size as higher risk thresholds are accepted. However, strong spatial variability in the optimal radius values across Europe supports the influence of landscape structure and environmental context (Figure 4). Smaller radii should be sufficient in heterogeneous areas (e.g., Bavarian Forest National Park), whereas larger radii would be required in more homogeneous areas (e.g., the Alps). This spatially explicit representation provides a practical tool for guiding the delineation of ASF restricted zones and can be readily adapted to any study area using our developed application: https://github.com/tkuerschner/RestrictionZoneTool.

### Comparison with buffers used during previous ASF outbreaks

By comparing our predicted buffer sizes with those applied in past ASF epidemics, we estimated an average optimal buffer size of 4.31 km in Belgium and 4 km in the Czech Republic based on a risk threshold of 0.05 and a 14-time window. Lowering the risk threshold to 0.01 increases the predicted average buffer size to 5.71 km in Belgium and 6 km in the Czech Republic. Alternatively, extending the time window to 21 days, e.g., including the incubation period, or the additional time before a carcass is likely to be discovered, further increases the predicted average buffer to 5.97 and 6 km for a 0.01 risk threshold, and 4.51 and 4.14 km for a 0.05 risk threshold, for Belgium and the Czech Republic, respectively.

## Discussion

Policy guidance for halting the spread of wildlife diseases typically specifies default restricted-zone sizes for rapid response (e.g., infected/ control/ surveillance zones). However, several studies show that the movement of wild animals and the landscape structure modulate disease dynamics (White, Forester & Craft 2018; Forero-Muñoz *et al*. 2025). This implies that standard restricted zones may be irrelevant if they do not take these factors into account. In this study, we modeled the risk that a wild boar leaves an area after entering it, across multiple spatial scales, to guide restricted zone definition for ASF management in wild boar populations. We found that the risk of leaving increased non-linearly over time and was strongly influenced by spatial scale and environmental conditions. Specifically, the risk of leaving an area decreased exponentially as its radius increased; however, homogeneous, hilly, or heavily human-modified landscapes increase the risk, especially at a fine spatial scale. This emphasizes the need to create adaptive, context-specific restricted zones. Beyond its applied relevance, these findings provide new insights into how temporal and spatial processes jointly shape movement decisions.

### Spatiotemporal dynamics of the risk of leaving

The observed “time-dependent risk” reflects a natural tendency for wild boar to disperse or expand movements to fulfill their requirements (Nathan *et al*. 2008; Morales *et al*. 2010; Jeltsch *et al*. 2013). However, this temporal pattern was not uniform, as the risk of leaving was strongly influenced by spatial scale and environmental conditions, highlighting that both habitat quality and duration of staying influence movement decisions. At larger spatial scales, the risk of leaving declined sharply and then plateaued, likely because broader spatial extents capture sufficient habitat heterogeneity to meet resource needs (Bauder *et al*. 2020). This plateau, along with the reduced effect of environmental covariates, is consistent with home range size approaching equilibrium (Saltz & Getz 2021; Wielgus *et al*. 2023b). At smaller scales, however, time and environmental conditions such as land cover composition and human impact strongly shaped movements, suggesting that wild boar respond mainly to the local environment.

We found that the risk of leaving increases with higher proportions of all land cover types, suggesting that wild boar is more likely to leave homogeneous areas dominated by a single land cover type. At larger spatial scales, heterogeneous environments are more prevalent, offering broader resource availability, more refuge options, and greater habitat diversity. This reduces the need for wild boar to disperse; such heterogeneous landscapes are generally characterized by smaller home ranges and lower movement rates (Walter *et al*. 2018; Bauder *et al*. 2020).

Human activities typically impact the movement of wild boar, resulting in increased movement, and reduced home range size (Podgórski *et al*. 2013; Johann *et al*. 2020; Davidson, Malkinson & Shanas 2022; Wielgus *et al*. 2024). We generally found similar patterns, with an increasing risk of leaving with human impact, as indicated by the proportion of cropland, the human modification index and the forest landscape integrity index (FLII). These patterns suggest that wild boar respond sensitively to immediate habitat quality and human presence at fine spatial scales. However, two unexpected patterns emerged: (1) At small spatial scales (≤ 2 km), higher FLII values were associated with a lower risk of leaving, consistent with the idea that intact, undisturbed forests provide shelter, refuge, and stable resources that encourage site fidelity; and (2) the influence of cropland persisted even at broader spatial scales. Although cropland provides an abundant food source (Herrero *et al*. 2006; Cappa, Lombardini & Meriggi 2019), its openness and associated hunting pressure may explain the consistently higher risk of leaving (Davidson, Malkinson & Shanas 2022). This suggests partial adaptation of wild boar to arable land.

Wild boar space use is fundamentally dynamic, but our findings emphasize that habitat composition and broader ecological and anthropogenic factors interact to influence movement decisions in wild boar. This pattern aligns with previous single-site studies showing that movement patterns and home range size of wild boar are shaped by multiple external factors, such as habitat quality, availability of food and shelter, topography, management practices (e.g., feeding, baiting) and human disturbance (such as hunting and urbanization; Massei *et al*. 1997; Truvé & Lemel 2003; Thurfjell, Spong & Ericsson 2013; Thurfjell, Spong & Ericsson 2014; Bisi *et al*. 2018; Amendolia *et al*. 2019).

### Implications for management of ASF in wild boar populations

Habitat heterogeneity should be considered when predicting wild boar movements and planning control measures. For example, heterogeneous environments may act as “retention areas”, where individuals are more likely to remain, whereas homogeneous areas tend to promote movement and dispersal. Management should therefore aim to maintain sufficient and diverse habitat, especially at the fine spatial scale, to promote site fidelity (Gardiner *et al*. 2019). In the context of ASF surveillance, we highlight that carcass searches and containment efforts should focus on heterogeneous landscapes, and particularly in forested areas, close to water courses and forest edges, where carcass deposition is most frequent (Morelle *et al*. 2019; Rogoll *et al*. 2024). In such environments, carcass removal can be carried out more efficiently and rapidly, enhancing the effectiveness of control efforts.

In general, a restricted zone with a radius of ∼8 km around an ASF-affected carcass would reduce the risk of disease spread by preventing infected wild boar from dispersing out of the control region within 14 days, regardless of environmental conditions. With an acceptable risk level of 5%, the width of the buffer could be reduced to 5 km in highly heterogeneous environments, even in areas of high human impact. Conversely, in homogeneous landscapes the buffers must be enlarged. Even a radius of 20 km may be insufficient when the proportion of agricultural land exceeds 75% of the observed area. However, it should be noted that this interpretation is specific to the 14-day risk of leaving and the 5% threshold; changing either the time window or the acceptable risk threshold alters the estimated buffer widths. In practice, these parameters depend on several factors. For instance, the time window should account for when the carcass was discovered, since older carcasses increase the likelihood of wild boar encountering them and becoming infectious themselves. Accurately determining the time of death is crucial to adapt the time window and ensure the buffer reflects the true period of risk. This can be based on factors such as the decay stage or ambient temperature (Müller *et al*. 2024). Additionally, the ASF virus spreads at a rate of 0.5-5 km per month, depending on the density of wild boar and human activity (Podgórski & Smietanka 2018; Vaclavek 2019; Taylor *et al*. 2021; Guberti *et al*. 2022). Therefore, the buffer should increase by 0.5-5 km for each month that the current outbreak is undetected.

Policymakers and stakeholders determine the acceptable threshold for the risk of leaving a buffer, and a cost-benefit analysis of the potential outcomes must be carried out (Rivas *et al*. 2012; EFSA Panel on Animal Health and Welfare *et al*. 2021; Lyons *et al*. 2021). However, the risk of spread from zones of different sizes must be quantified as a prerequisite. The interactive tool that we have developed allows users to generate maps of the optimal radius around areas affected by ASF, based on adjustable risk thresholds and time windows. The risk threshold that has been chosen is the proportion of wild boar that we are ‘willing to accept’ may leave the specified area. This approach highlights the dynamic nature of wild boar movement and disease spread, demonstrating that optimal control measures depend on the context. By allowing stakeholders to explore how different thresholds and temporal scales influence buffer size, the tool provides a flexible decision support system that can be adapted to local ecological and management conditions. We anticipate that this work and the associated user-friendly tool could aid the management of future ASF outbreaks in wild boar populations.

Our predicted optimal buffer sizes were broadly comparable to those implemented in past ASF epidemics, but they also provide a more quantitative, risk-based rationale for their selection. For instance, for a 14-day period and a 0.01 risk threshold, predicted buffers were 5.71 km in Belgium and 6 km in Czech Republic, similar to the buffers used to determine the infected area and fenced off (6 km, More *et al*. 2018; OIE 2019; Šatrán 2019; Jori *et al*. 2021; Smith *et al*. 2022). Longer time windows (21 days) led to larger predicted buffers, suggesting that previously applied zones may have underestimated the area needed to contain wild boar movements in certain contexts. This comparison demonstrates that our proposed buffer sizes are realistic and conservative in relation to known disease spread dynamics. It also illustrates that our approach could complement existing guidelines by providing site-specific, biologically informed buffer sizes, potentially improving the effectiveness and efficiency of ASF containment measures.

It is worth mentioning that our results are likely to be a conservative estimate of the buffer size. Sick animals, including ASF-infected wild boar, tend to move less (Morelle *et al*. 2023; Grabow *et al*. 2024; Grabow *et al*. 2025), but obtaining movement data from ASF-infected wild boar in free-ranging populations is obviously difficult. Here, we especially capture individuals that are newly infected by contact with a sick animal, given that the incubation period is about one week. To adopt a more conservative approach, we also provide a three-week buffer estimate, accounting for one week of incubation from a freshly deceased animal plus the additional two-week critical time window.

### Methodological contributions

While a previous work by Wielgus *et al*. (2023a) demonstrated the value of an integrated FPT-survival analysis approach for risk of leaving, the novelty here lies in the incorporation of environmental factors. By considering habitat characteristics, landscape features, and human influence, the method allows us to examine how movement risk changes with spatial scale and environmental conditions. This represents a significant improvement over the standard uniform approaches typically applied in disease management, moving towards a more ecologically informed and spatially adaptive delineation of infected areas.

A key aspect of our work was emphasizing the importance of examining animal spatial behavior over short, disease-relevant timeframes (i.e., 14 days). Such timescales are often overlooked in wildlife ecology, where the focus tends to be on longer-term movement patterns (Kay *et al*. 2017; Mumme *et al*. 2023, E. Wielgus, unpublished data). By aligning risk estimation with disease progression and transmission windows, and by integrating environmental conditions, we demonstrate how movement risk can be quantified in ways that are both ecologically and epidemiologically meaningful. The method has considerable potential for application to future outbreaks of other infectious diseases beyond ASF. The framework is also highly flexible and can be adapted to assess movement risk at different temporal and spatial scales, depending on the disease context and management needs.

## Acknowledgements

This work was carried out with the support of the EUROMAMMALS/EUROBOAR collaborative project (https://euromammals.org/euroboar/). We thank all the project members who supported and contributed to this initiative, as well as the many people across Europe who helped to capture and tag the animals studied in this article.

