## Supplementary Information for "How big is enough? Movement-informed zoning for African swine fever mitigation"

1. Correlation between predictor variables

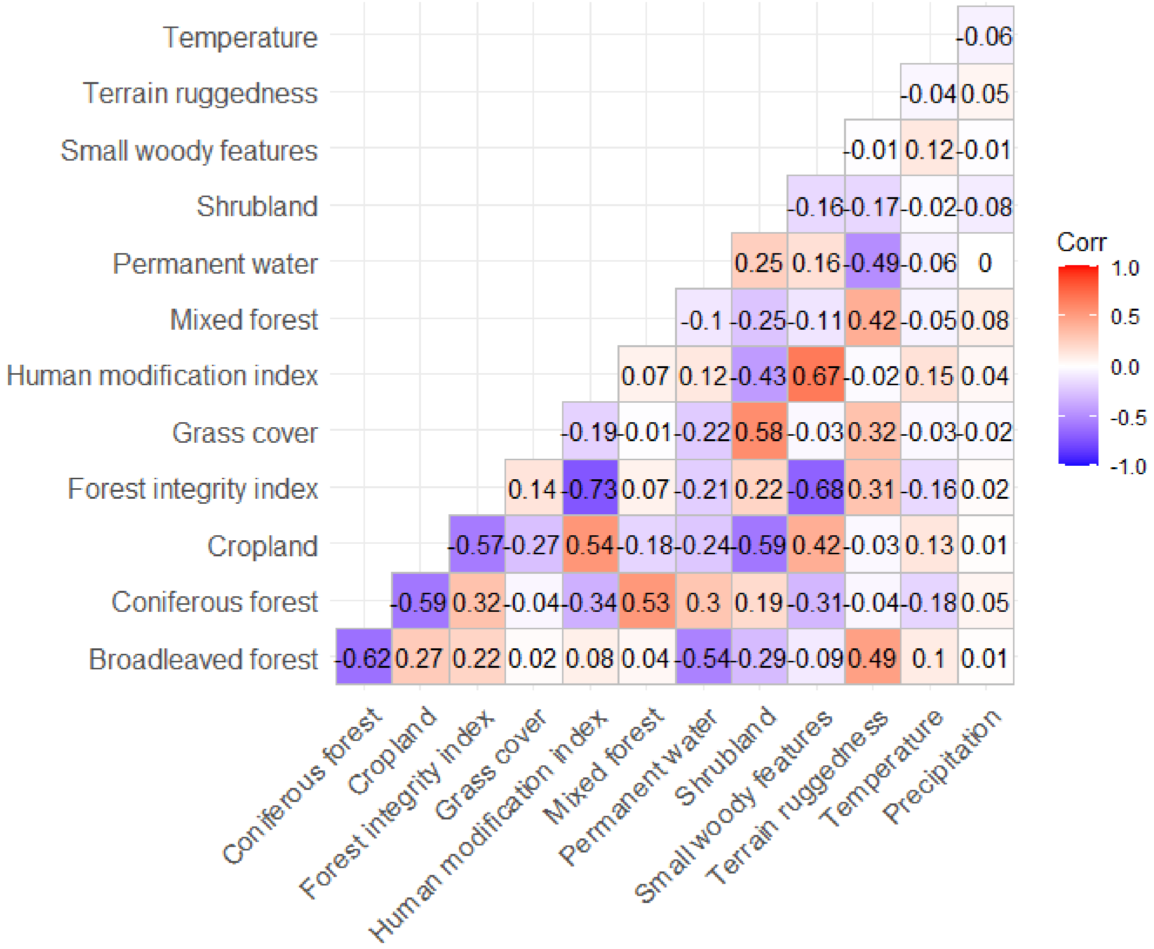


**Figure S1.** Correlation matrix of predictor variables included in the Cox proportional hazards model. Each cell represents the pairwise Spearman correlation coefficient between two variables. Blue colors indicate negative correlations, red colors indicate positive correlations, and the intensity of the color corresponds to the strength of the correlation.

### 2. Results CPH models

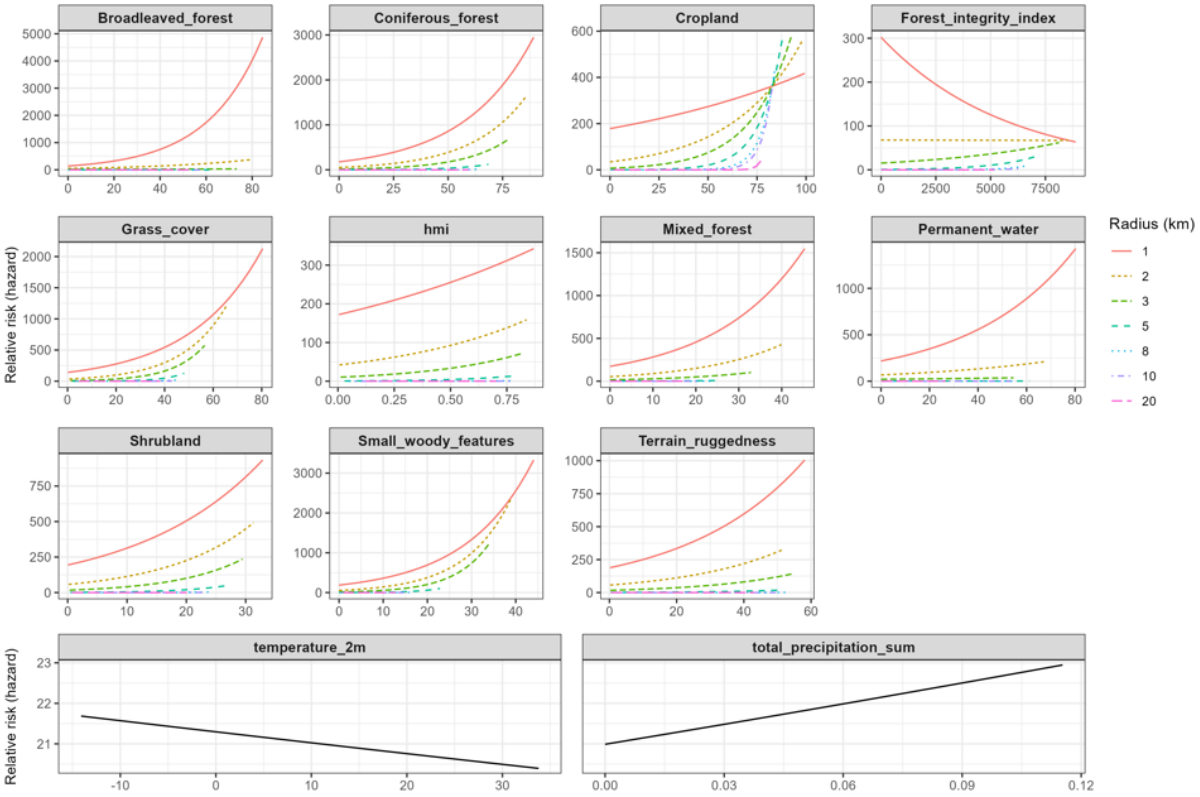


**Figure S2.** Relative risk hazard (i.e., at any given time)

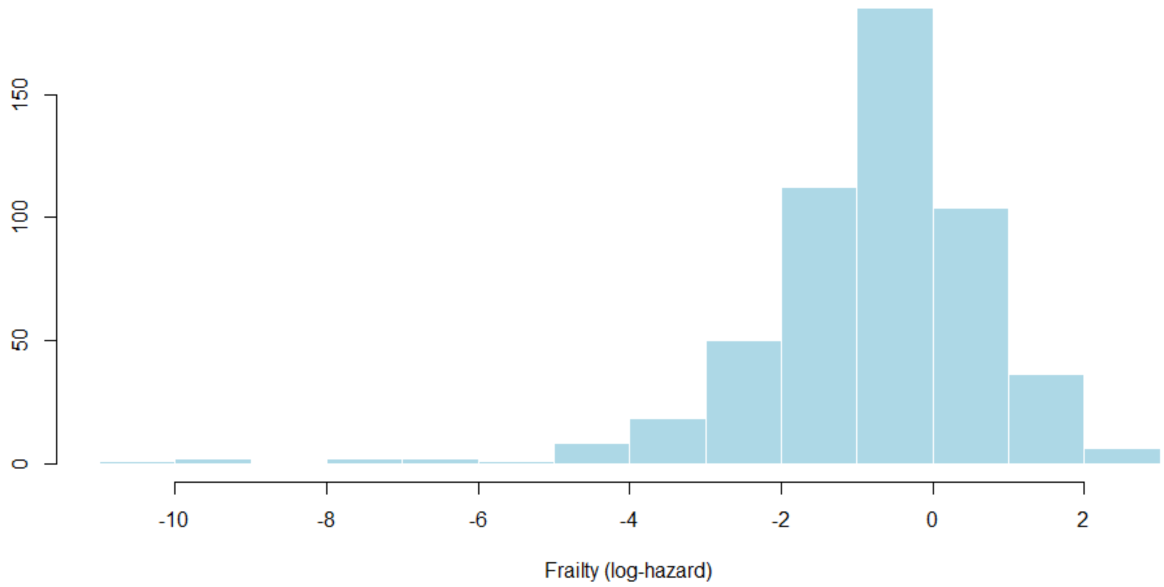


**Figure S3.** Distribution of individual frailty effects estimated from the Cox proportional hazards model. Positive values indicate higher-than-average hazard (i.e., higher risk than population average), while negative values indicate lower-than-average. The spread of the distribution reflects variability in baseline risk across individuals.

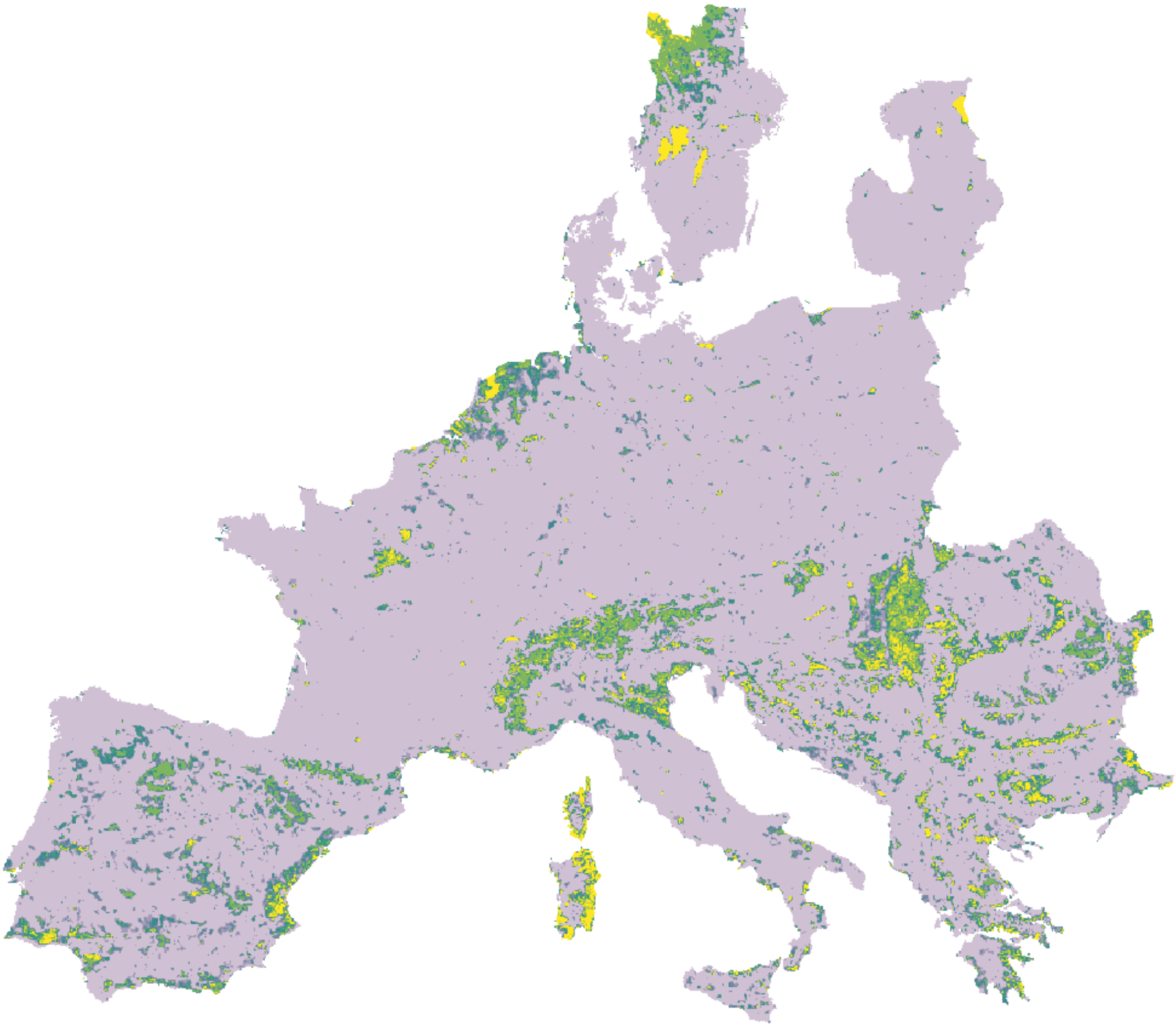

**Figure S4.** Extrapolation map. Green to yellow indicates extrapolation in 50% to 100% of the radius size.
